# Dopamine maintains network synchrony via direct modulation of gap junctions in the crustacean cardiac ganglion

**DOI:** 10.1101/355651

**Authors:** Brian J Lane, Daniel R Kick, David K Wilson, Satish S Nair, David J Schulz

## Abstract

Abstract The Large Cell (LC) motor neurons of the crab (C. borealis) cardiac ganglion have variable membrane conductance magnitudes even within the same individual, yet produce identical synchronized activity in the intact network. In our previous study (*Lane et al., 2016*) we blocked a subset of K^+^ conductances across LCs, resulting in loss of synchronous activity. In this study, we hypothesized that this same variability of conductances could make LCs vulnerable to desynchronization during neuromodulation. We exposed the LCs to serotonin (5HT) and dopamine (DA) while recording simultaneously from multiple LCs. Both amines had distinct excitatory effects on LC output, but only 5HT caused desynchronized output. We further determined that DA rapidly increased gap junctional conductance. Co-application of both amines induced 5HT-like output, but waveforms remained synchronized. Furthermore, DA prevented desynchronization induced by the K^+^ channel blocker tetraethylammonium (TEA), suggesting that dopaminergic modulation of electrical coupling plays a protective role in maintaining network synchrony.

## Introduction

Neural networks must be capable of producing output that is robust and reliable, yet also flexible enough to meet changing environmental demands. One mechanism of providing flexibility to network activity is neuromodulation, which reconfigures network output by altering a subset of cellular and synaptic conductances (*Harris-Warrick, 2011; Bargmann, 2012; Daur et al., 2016*). However, many networks achieve stable output by a variety of solutions; intrinsic membrane conductances and synaptic strengths can be highly variable yet still produce nearly identical physiological activity (*Ball et al., 2010; Calabrese et al., 2011; Marder, 2011; Ransdell et al., 2013b*). This raises a fundamental question about neuromodulation, highlighted in a recent review by *Marder et al. (2014),* as to whether modulation of networks with variable underlying parameters can produce predictable and reliable results. These authors demonstrate computationally that modulation of neurons with similar outputs arising from variable underlying conductances can cause anywhere from relatively small to fairly substantial differences in output (*Marder et al., 2014*). Therefore, the response of any neural network to modulation is likely state-dependent (*Goldman et al., 2001; Nadim et al., 2008; Gutierrez and Marder, 2014; Williams et al., 2013; Marder et al., 2014*), and potentially unpredictable as a result of these varying underlying conductances (*Marder et al., 2014*). In some cases neuromodulation can expand the parameter space in which a given activity feature is maintained (*Grashow et al., 2009*), potentially leading to protective effects of modulation that ensure robust network output (*Städele et al., 2015*). Yet these questions have never been addressed in a network that relies on synchronous activity for appropriate physiological output.

The crustacean cardiac ganglion (CG) is a central pattern generator network that produces rhythmic bursts with precisely synchronized activity across all five Large Cell (LC) motor neurons (*Lane et al., 2016*), constituent LCs are variable in many ionic conductances, including A-type K+ (IA), high-threshold K+ (IHTK), and voltage-dependent Ca2+ (ICa) (*Ransdell et al., 2013a*, b; *Lane et al., 2016*). When subsets of K+ conductances are blocked in LCs of a network, synchrony is disrupted, although ultimately is restored by a combination compensatory changes in membrane conductance and electrical synaptic strength (*Lane et al., 2016*). As the CG is modulated by many substances, including neuropeptides and the biogenic amines serotonin and dopamine (*Cooke, 2002; Cruz-Bermudez and Marder, 2007*) that are known to target the same K+ conductances that lead to desynchronization when altered (*Kloppenburg et al., 1999; Peck et al., 2001; Johnson et al., 2003; Gruhn et al., 2005*), then one potentially detrimental impact of neuromodulation is the resulting loss of LC synchrony. Given that hormonal modulators in the hemolymph will bathe LCs uniformly, this study addresses whether the CG is tuned to maintain stable synchrony during neuromodulation or if altering a subset of cellular conductances with neuromodulation will desynchronize network activity.

We hypothesized that neuromodulation will desynchronize LC activity owing to the variable conductances across LCs. We tested this hypothesis by exposing the CG to two amine modulators, serotonin (5HT) and dopamine (DA), and measuring the effects of the modulators on excitability and synchrony individually and in conjunction. We found serotonergic modulation desynchronizes LC voltage waveforms, and in most networks elicited prolonged pacemaker bursts driving two distinct bursts in LCs before the cycle was reset. In contrast, DA had an overall excitatory effect on LCs, but neither desynchronized LC activity nor elicited the burst doublets seen in 5HT. When co-applied, DA prevented the 5HT-induced desynchronization without preventing the 5HT output in the form of characteristic burst doublets. Even with IHTK reduced by TEA application - a known perturbation that leads to substantial loss of synchrony across LCs (*Lane et al., 2016*) - co-application of DA with TEA prevented desynchronization. Our results suggest that DA preserves network synchrony by directly targeting and increasing electrical synaptic conductance. Thus DA may function to maintain robust synchrony in the cardiac network while still being permissive to plasticity of output caused by other modulators.

## Results

### 5HT and DA have distinct excitatory effects when applied to the entire network

Both 5HT (10^−6^M) and DA (10^−5^M) are excitatory when applied to the entire CG of C. borealis (*Cruz-Bermudez and Marder, 2007*), and our results recapitulate this same effect (Figure 2). 5HT significantly increased pacemaker burst duration and in 6 out of 8 experiments switched the network to a distinct output consisting of a single prolonged pacemaker burst driving two distinct LC bursts (Figure 2A). Changes in network output in 5HT were quantified by statistical analysis of LC spiking from extracellular recordings (Figure 2B, N=8). 5HT significantly decreased interburst interval (4.15±3.07 in control; 2.04±0.46 in 5HT; P<0.05) and cycle period (5.149±3.086 in control; 2.607±0.563 in 5HT; P<0.05). 5HT increased the number of spikes per burst (6.250±1.883 in control and 9.625±2.789 in 5HT; P<0.01), the spike frequency within each burst (5.640±3.995 in control and 16.133±3.417 in 5HT; P<0.01), and LC duty cycle (0.161±0.0799 in control and 0.241±0.0530 in 5HT; P<0.05).

DA also increased network excitability, but in distinct ways from 5HT (Figure 2C). DA did not induce doublet bursting, but significantly increased the number of LC spikes per burst (8.850±3.978 in control; 11.225±5.337 in DA; P<0.05), LC spike frequency (8.854±4.878 in control; 11.268±7.434 in DA; P<0.05), LC burst duration (0.715±0.280 in control; 0.806±0.281 in DA; P<0.05), and LC duty cycle (0.164±0.0744 in control; 0.211±0.0411 in DA; P<0.05) (Figure 2D, N=8).

**Figure 1.**
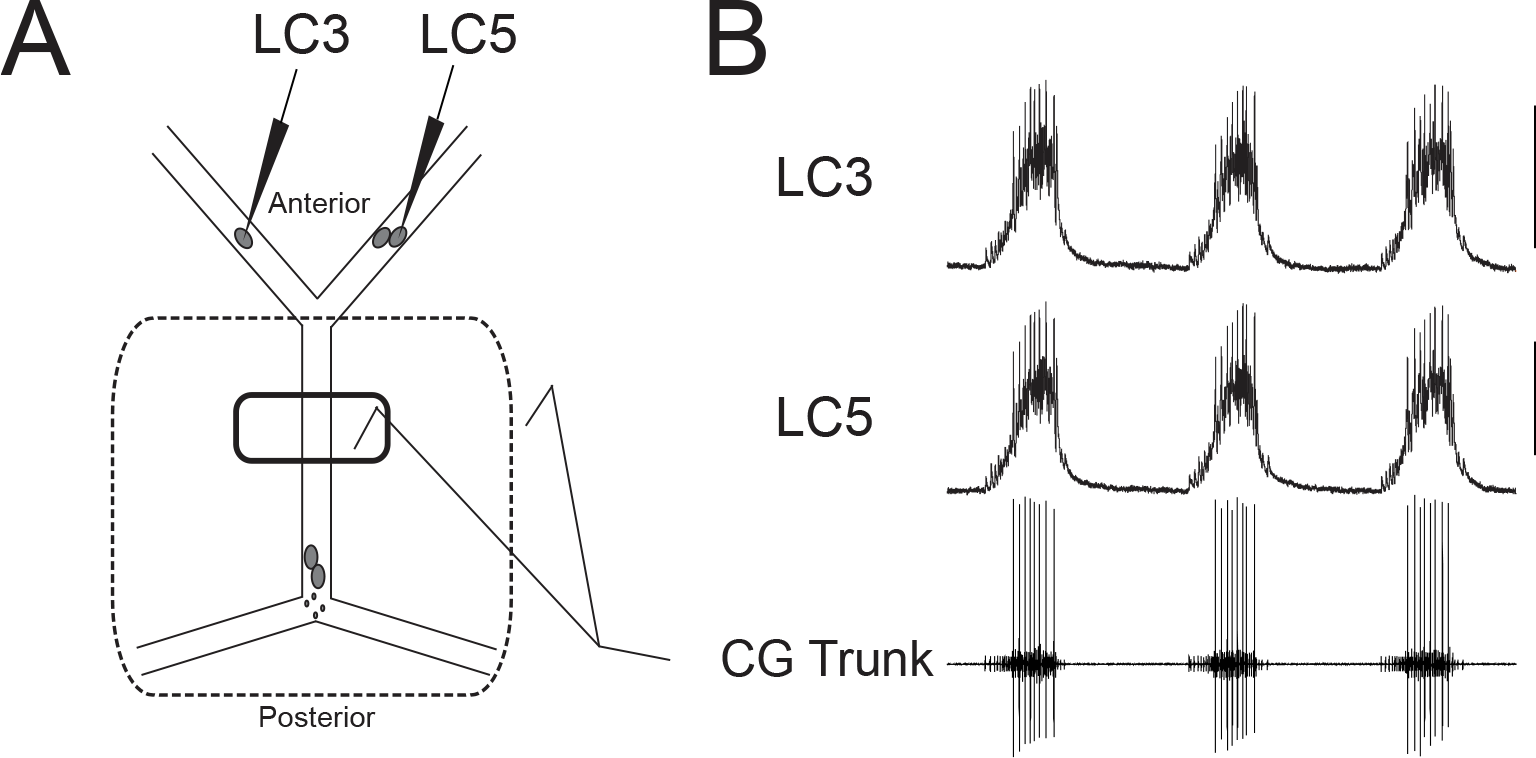
Experimental Setup and Typical Activity in the CG. **A.** Intracellular electrodes recorded simultaneously from LC3 and either LC4 or LC5. Extracellular recordings were taken from a petroleum jelly well on the CG trunk (connecting line). For experiments which applied modulators or TEA exclusively to anterior LCs, the extracellular recording was taken from a single large well allowing anterior LCs to be exposed to the perfusate while protecting the remainder of the ganglion (dashed line). **B.** Representative control activity showing the rhythmic synchronized bursting of LCs. (Scale bars = 10mV, recording length = 14 sec).

When applied focally to the anterior LCs at these same concentrations, our results demonstrate that both amines have direct excitatory effects on LCs. 5HT applied only to the anterior LCs increased the number of spikes per burst (5.850 ±2.72 in control; 10.85 ± 2.96 in 5HT; P<0.01), the spike frequency within each burst (6.24 ±3.18 Hz in control; 11.53 ± 3.39 Hz in 5HT; P<0.01), burst duration (0.642 ± 0.367 sec in control; 0.739 ± 0.411 sec in 5HT; P<0.05), LC duty cycle (0.183 ±0.059 in control; 0.232 ±0.063 in 5HT; P<0.01) and decreased the interburst interval (3.15±0.68 in control; 2.761 ±0.658 in 5HT; P<0.01). DA also directly affected LC output, significantly increasing the number of spikes per burst (4.45 ±3.42 in control and 10.20 ±6.41 in DA; P<0.05) and spike frequency within each burst (5.85 ±4.95 in control and 10.38 ±10.16 in DA; P<0.05).

### 5HT Desynchronizes Burst Waveforms but DA Does Not

The degree of synchrony in burst waveforms was quantified as described previously (*Lane et al. (2016*), see Methods). Briefly, we performed a cross-correlation on the digitized voltage waveforms of each burst from two intracellular recordings. The coefficient of determination (R^2^) was then used to quantify how accurately the voltage of one cell predicts the voltage of the other. This provides a baseline measure of synchrony for each burst and allowed us to track relative changes.

At the onset of serotonergic modulation (10^−6^M), there was an acute reduction in synchrony as measured by R^2^. Figure 3A illustrates typical acute effects of 5HT application. Differences in voltage waveforms appear between LC3 and LC5 after 5HT perfusion. R^2^ values for every burst during a single preparation are plotted in Figure 3B to visualize the changes in synchrony evident during these experiments. Synchrony reliably reached a minimum within several minutes (mean = 9.1 minutes) before stabilizing and showing a slow, modest recovery. For statistical analysis in Figure 3C, R^2^ values from 10 consecutive bursts were averaged at 3 time points: Control (5 minutes prior to modulation), acute 5HT modulation (sampled at the point of maximum desynchronization), and again after 30 minutes of exposure to 5HT (N=8 preparations). Synchrony significantly decreased in 5HT (R^2^ = 0.967±0.012 control, 0.893±0.081 5HT; p<0.01, Signed rank test). Subsequently there was a significant increase in synchrony between acute modulation and 30 minutes of continuous 5HT perfusion (0.927±0.066) (p<0.01, Signed rank test), but the cause of this increase is not clear. However, synchrony was still significantly below baseline after 30 minutes (p<0.05, Signed rank test).

**Figure 2.**
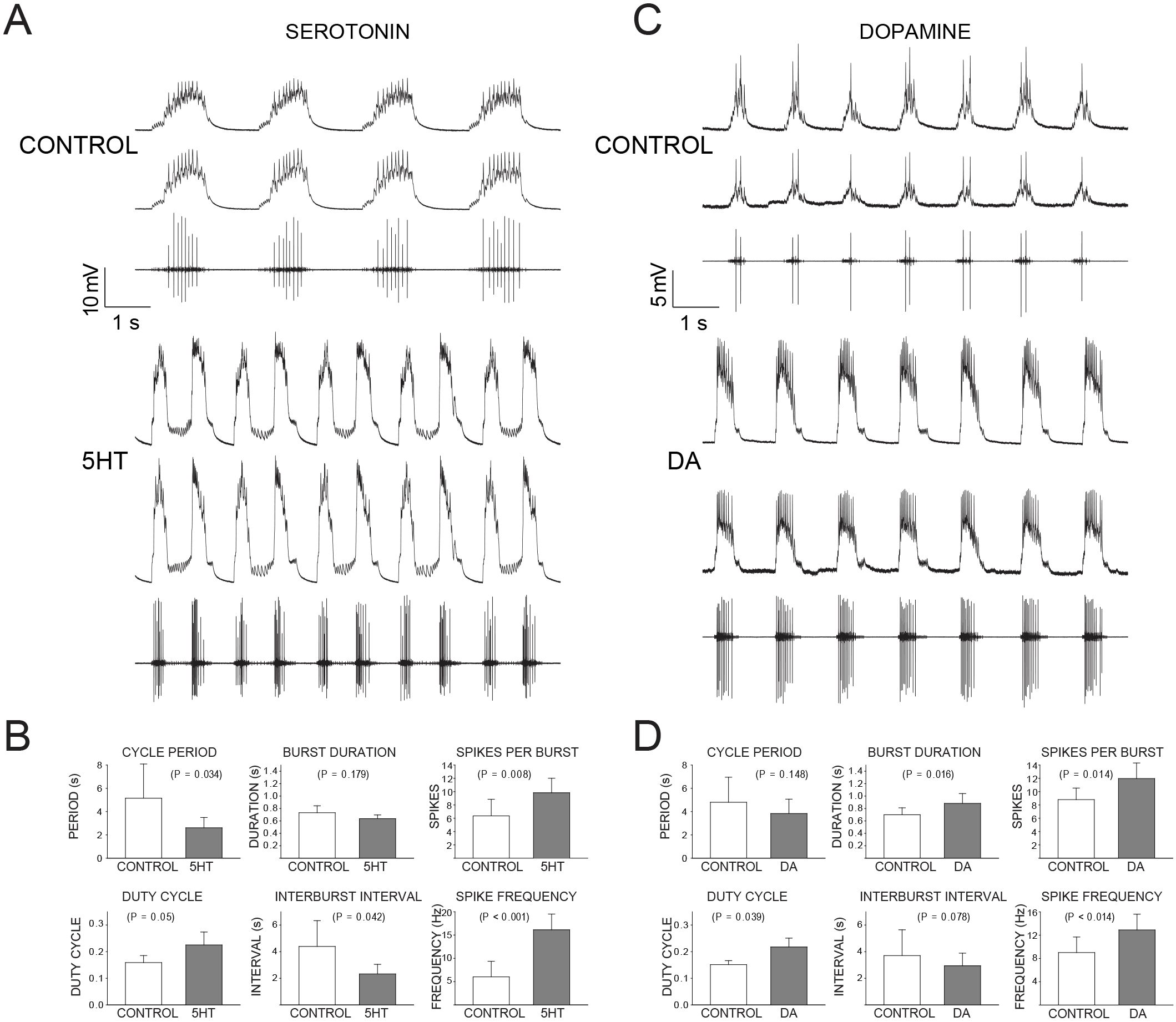
Effects of 5HT and DA on output of the CG network. **A.** Effects of 5HT on the cardiac network. For each condition (Control and 5HT) the top two traces are intracellular recordings from two different LCs in the same network (LC3 and LC4) and the bottom trace is an extracellular recording from the CG trunk. 5HT was applied at 10^−6^M. **B.** Quantitation of burst properties in Control and 5HT CG networks. Data were taken from LC spikes on the extracellular recordings. Bars represent mean +/− SD. P-values from comparisons of Control vs. 5HT via paired t-tests are reported for each measurement. **C.** Effects of 10^−5^M DA on the cardiac network. Recordings as in panel A. **D.** Quantitation of the effects of DA on LC burst output, as described in panel B.

During 5HT-induced doublets, the first and second of the two bursts typically displayed distinct waveform synchrony values. This can be seen from between 10-30 minutes for the preparation shown in Figure 3B, in which R^2^ values alternated between the upper and lower bands. The R^2^ values for both the 1st and 2nd burst were below control (p<0.01, signed rank test), but which of the two bursts in the doublet had lower R^2^ values varied across preparations.

In stark contrast, burst waveforms for LCs in DA remained completely and strikingly synchronized (Figure 3D). The scatterplot in Figure 3E shows R^2^ for each burst in one preparation during a full 30 minutes of DA perfusion. There were no noticeable changes in synchrony during any DA perfusion experiment (Figure 3E; 3F): there were no significant changes in R^2^ from control (0.964±0.026), acute DA modulation (0.966±0.025), and 30 minutes of DA modulation (0.966±0.024). The acute data point was sampled at 9.1 minutes to match the mean time point used for sampling in 5HT.

### DA, but not 5HT, modulates coupling conductance

We hypothesized that modulation of electrical coupling may be responsible for the different effects of 5HT and DA on LC synchrony. One possibility is that modulation with 5HT does not increase the strength of electrical coupling sufficiently to prevent differential effects on cell excitability from resulting in desynchronization. DA may increase coupling strength enough to ensure synchrony.

As an indicator of the strength of electrical coupling, we measured the coupling coefficient (see Methods) between LC3 and LC5 at control and after 15 minutes of modulation. Coupling coefficients were not significantly different between control (0.043± 0.035) and 15 minutes modulation in 5HT (0.037 ± 0.042) (Figure 4A; N=8). In contrast, DA increased coupling coefficient by 41 percent (0.035±0.030 in control; 0.049±0.043 in DA) (P<0.05, N=6; Figure 4A). Input resistance was not significantly changed in either 5HT (3.62±4.40 megaOhm in control; 3.78± 4.11 megaOhm in 5HT) or DA (3.85±2.34 megaOhm in control; 3.33±1.77 megaOhm in DA) (Figure 4A). These results suggest that an increase in coupling coefficient in DA may be the result of a direct increase of coupling conductance in the presence of DA.

While the coupling coefficient is a useful description of the functional coupling relationship, it does not identify the electrophysiological mechanism, i.e. a change in membrane resistance or a change in coupling conductance. The branch containing LC4 and LC5 can be isolated from the network by thread ligature creating the ideal conditions for measuring the coupling conductance between these somata (Figure 4B). With two electrodes in each cell, we used hyperpolarizing current injections to determine the gap junctional resistance independent of membrane resistance (see Methods). 5HT had no apparent effect on coupling conductance, while DA significantly increased coupling conductance in both directions between LCs 4 and 5 within 15 minutes of DA exposure (Figure 4C; 4D).

### Coapplication of DA and 5HT Prevents Desynchronization and Induces Doublet Bursting

Our previous work demonstrated that a compensatory increase in electrical coupling among LCs rescued synchrony after treatment with the channel blocker TEA (Lane et al. 2016). The relatively rapid and large increase in electrical coupling between LCs upon exposure to DA led us to hypothesize that co-modulation with DA might prevent desynchronization caused by other modulators. We tested the ability of DA to maintain network synchrony during co-modulation with 5HT using the same perfusion and recording protocol as above, by co-applying DA (10^−5^ M) and 5HT (10^−6^ M).

Individual preparations treated with DA+5HT sometimes showed a small increase or transient decrease in synchrony, which occurred over the same time scale as seen in preparations exposed to 5HT alone. However, across the full set of preparations (N=8) there was no significant change in synchrony when DA and 5HT were co-applied. 5 out of 8 preparations transitioned to the doublet bursting mode seen in 5HT alone (Figure 2A, 5A). However, this time the double-bursting pattern displayed highly synchronized waveforms (Figure 5A). The scatterplot in Figure 5B shows R^2^ for all bursts across a full experiment, in this case revealing a slight increase in synchrony from baseline after the onset of co-modulation and, in contrast with Figure 2B, no clear segregation of the R^2^ values for the 1st and 2nd burst in each doublet. Overall, there was no significant difference in synchrony of control (R^2^ = 0.971±0.01) and acute co-modulated preparations (R^2^ = 0.969±0.01; Figure 5C). However, there was a significant increase in synchrony between acute and 30 minutes (R^2^ = 0.983±0.006; P<0.01), suggesting a longer-term enhancement of coupling throughout the experiment.

**Figure 3.**
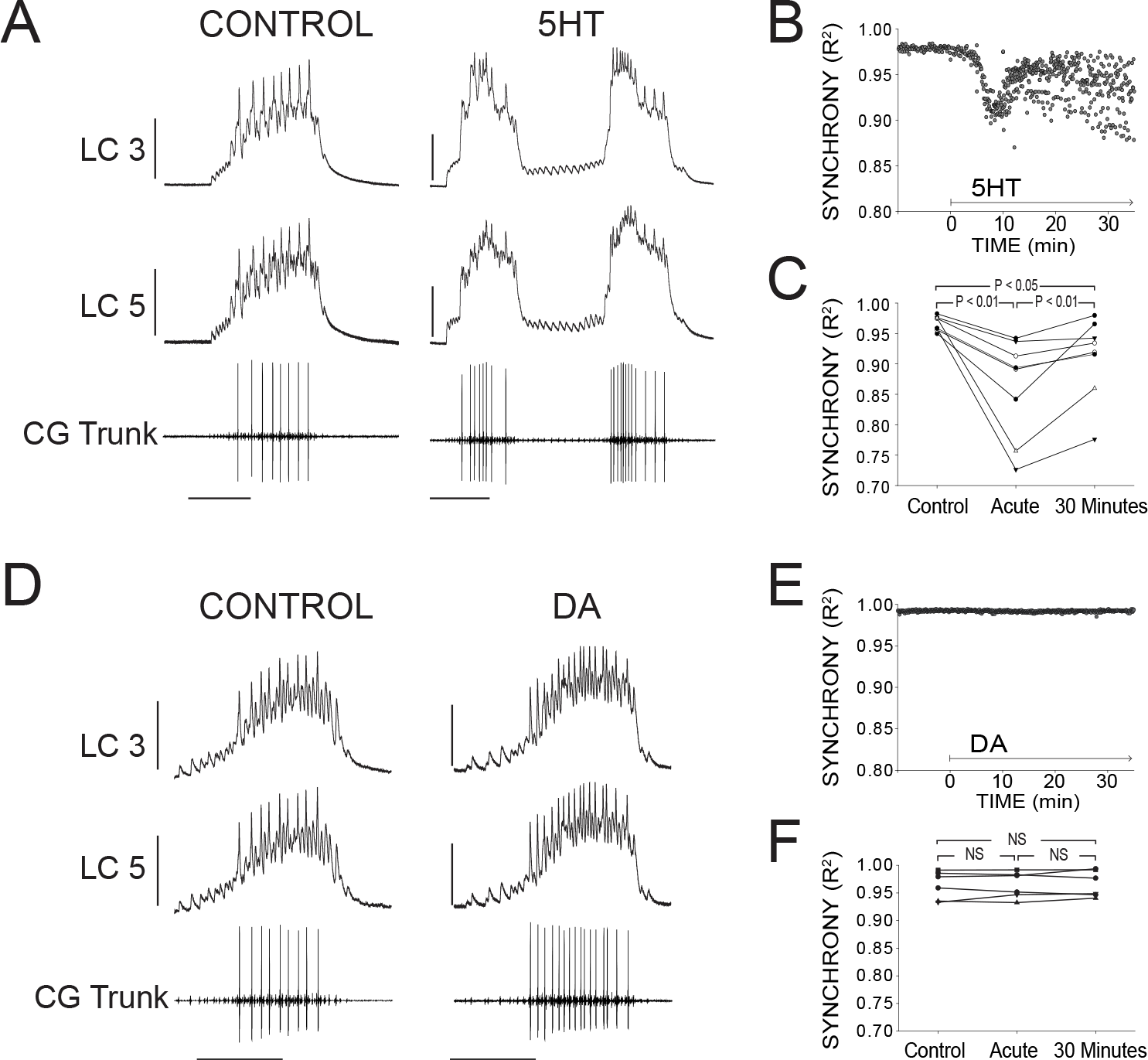
Effects of 5HT and DA on Synchrony of LC Voltage Waveforms. **A.** Representative traces show that LCs with virtually identical control activity produce different burst waveforms after application of 5HT. Bursts in 5HT occurred in doublets in most preparations (6 of 8). Scale bars = 10 mV and 1 second. **B.** Waveform synchrony (R^2^) was calculated for every burst across a full experiment. Scatterplot shows consistent synchronized bursting across 10 minutes of control activity followed by 30 minutes of continuous perfusion of 5HT. An acute loss of synchrony accompanies the onset of modulation. Doublet bursting results in two distinct bands of R^2^ values during 30 minutes of 5HT perfusion. **C.** R^2^ was averaged for 10 consecutive bursts at each of 3 time points: control (5 minutes prior to perfusion), Acute (at the point R^2^ reached a minimum), and after 30 minutes of modulation. A significant decrease occurred from control to acute (Signed Rank test, p<0.01), and a significant increase between acute and 30 minutes (Signed Rank test, p<0.05). Synchrony was not restored to control levels after 30 minutes (paired t-test, p< 0.05). N=8 preparations. **D.** Representative traces show that excitability and network output are affected by DA, but LCs remain synchronized. Scale bars = 10 mV and 1 second. **E.** R^2^ was calculated for every burst across a full experiment. Scatterplot shows this for 10 minutes of control activity followed by 30 minutes in DA. R^2^ values are largely unaffected by changes in activity caused by DA. **F.** R^2^ was averaged for 10 consecutive bursts at each of3 time points: control (5 minutes prior to perfusion), Acute (sampled from the same time-point as 5HT), and after 30 minutes of modulation. There were no significant differences between any two groups (paired t-tests). N=6 preparations.

**Figure 4.**
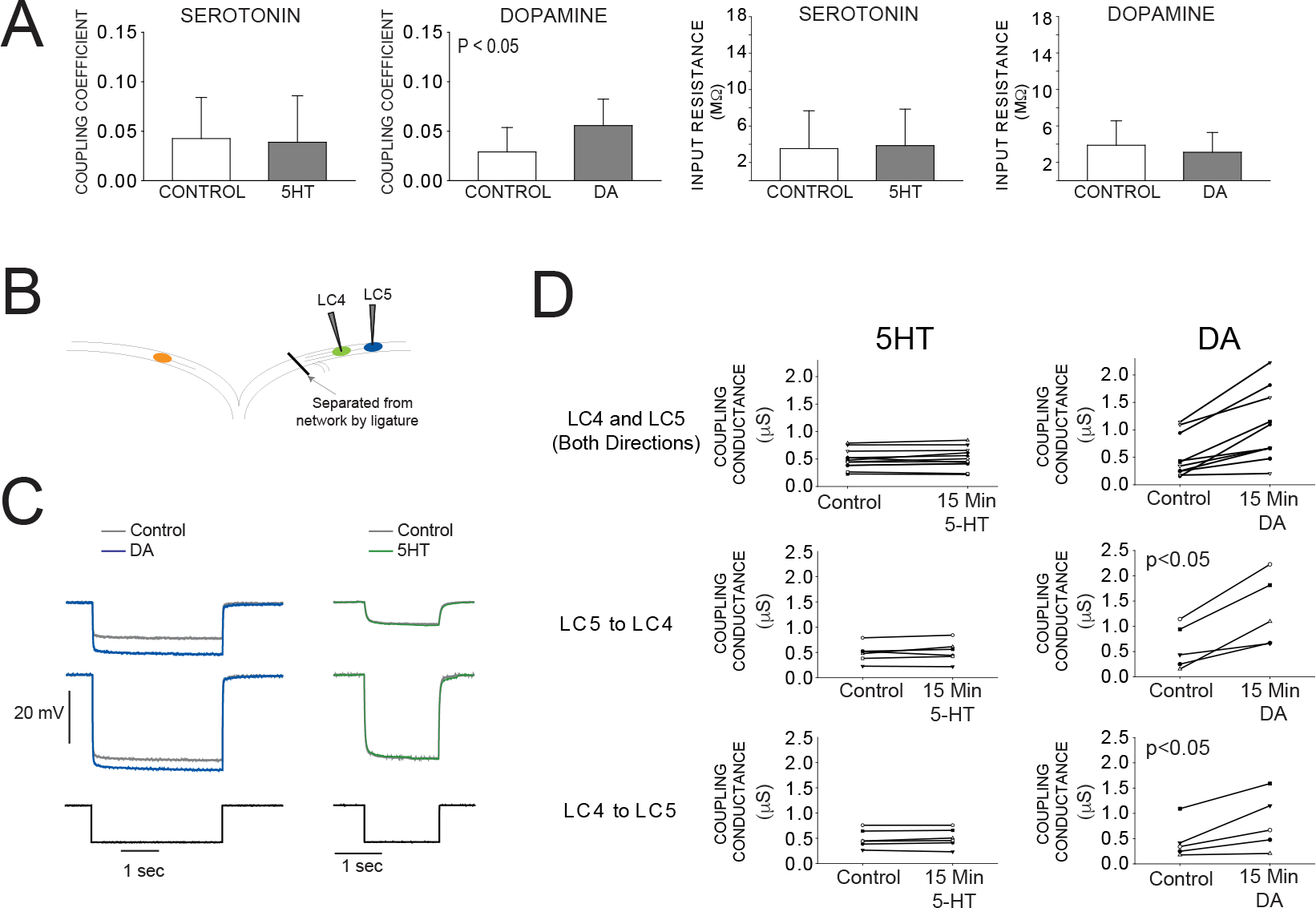
Effects of 5HT and DA on Electrical Coupling. **A.**Coupling coefficients for LC3-LC4 or LC3-LC5 in the active network were not significantly changed by 5HT. Mean coupling coefficient increased by 41 percent in DA (p<0.01). **B.** Diagram of reduced preparation used to test the effect of 5HT and DA on coupling conductance. LC4 and LC5 somata were physically isolated by thread ligature. **C.** Representative traces of current injections used to calculate coupling conductance before and after DA exposure (left) and 5HT exposure (right). Traces in grey show voltage responses of LC4 (top) and LC5 (middle) to a −8 nA hyperpolarizing current injection (bottom). Overlaid traces show the voltage response to the same current injection after DA (left, blue) and 5HT (right, green). **D.** Coupling conductance between LC4 and LC5 was unchanged by 5HT, and increased by DA. Top row in C shows coupling conductance in both directions for N=6 preparations before and after modulation. These data are then separated by directionality. Coupling Conductance was significantly increased in DA in both directions (p<0.05 for each, mean increase 149 percent, N=5).

Overall, DA+5HT co-modulation was strongly excitatory. Characteristics of LC output were affected by co-modulation (Figure 5D) similar to those found in 5HT alone (Figure 2B), including significant changes in the number of spikes per burst (2.600±4.034 in control and 11.53 ±3.58 in DA+5HT; P<0.001), spike frequency (4.840±3.818 in control and 10.345±3.144 in DA+5HT; P=0.05), burst duration (0.403±0.361 in control and 1.074±0.447 in DA+5HT; P<0.05), interburst interval (3.829±3.321 in control and 2.779±1.816 in DA+5HT; P<0.05), and LC duty cycle (0.0953±0.0425 in control and 0.275±0.0831 in DA+5HT; P<0.01).

### DA Prevents TEA Induced Desynchronization

We hypothesize that DA may prevent desynchronization against a variety of perturbations by modulating electrical coupling strength. Our previous study showed that exposing the anterior LCs to the K+ channel blocker TEA produces a substantial loss of LC synchrony as well as a large increase in the number of spikes per burst and spike frequency (*Lane et al., 2016*). For reference, the inset in Figure 6A (acute TEA, no DA) includes representative traces of TEA-induced desynchronization.

If DA can act broadly to maintain synchrony, then we hypothesized that DA should prevent desynchronization in TEA. To test this, a barrier of petroleum jelly was built to protect the posterior (pacemaking) end of the ganglion from the perfusate while leaving anterior LCs exposed (see Figure 1A). The anterior LCs were pre-incubated with DA for 5 minutes, and then the perfusion switched to saline containing both DA and TEA. The preparations pre-incubated in DA for 5 minutes did not show any loss of synchrony as a result of TEA (Figure 6B, 6C). Sample traces (Figure 6A) include 4 time points to show R^2^ at control (0.950±0.0149), 5 min DA (0.958±0.010), acute exposure to DA+TEA (0.953±0.011), and 30 minutes exposure to DA+TEA (0.968±0.008). Excitability increased after 5 minutes with the addition of DA, similar to those found after 15 minutes of DA in previous experiments (Figure 2D). After the addition of TEA at 5 minutes, there was a further increase in the number of spikes per burst (7.89±5.34 in DA and 16.39±7.65 in DA+TEA; P<0.01), spike frequency (8.25±7.95 in DA and 15.19±8.06 in DA+TEA; P<0.01), and cycle period (3.34±0.77 in DA and 3.97±1.06 in DA+TEA; P<0.01).

## Discussion

Neuromodulation represents a critical mechanism underlying circuit plasticity: by targeting subsets of ionic and synaptic conductances, modulators can reconfigure networks based on environmental feedback to produce entirely distinct circuit outputs leading to behaviorally relevant output that is adapted to the conditions at large (*Harris-Warrick, 2011; Nadim and Bucher, 2014*). However, a more recent appreciation of the inherent variability in underlying conductances of individual neurons, even within the same network of the same individual, poses a potential complication for the role of modulation in modifying circuit output. That is, how is reliable neuromodulation (and robust output) achieved when a common modulator targets variable neurons (*Grashow et al., 2009; Marder et al., 2014*)? Our study sheds some light on this potential conundrum by revealing complementary roles of the co-modulators 5HT and DA in changing circuit output yet ensuring some features of inherent circuit stability. Specifically, co-application of both modulators leads to a modification of circuit output similar to the effects of 5HT alone, but preserves the synchrony among individual neurons likely through the direct actions of DA on electrical synapses.

**Figure 5.**
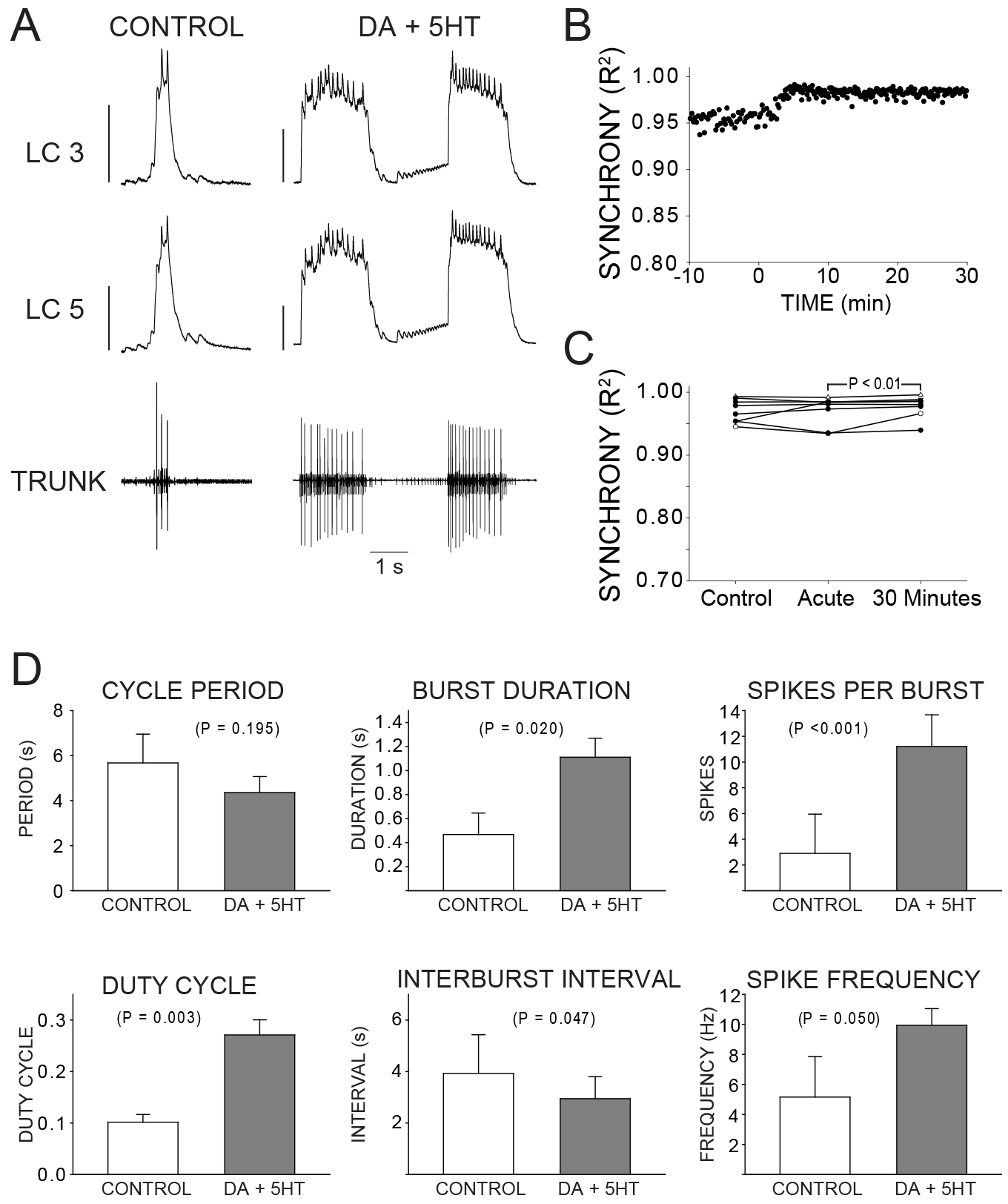
Effects of co-application of DA and 5HT on bursting output and synchrony. **A.** Representative traces show LCs maintain synchronized voltage waveforms during co-application of DA and 5HT. 5 out of 8 preparations transitioned to bursting in doublets, and network output shows increased number of spikes per burst, spike frequency in each burst, burst duration and LC duty cycle. Scale bars = 10 mV and 1 second. **B.** R^2^ was calculated for every burst across a full experiment. Scatterplot shows this for 10 minutes of control activity followed by 30 minutes of perfusion with both modulators. **C.** R^2^ was averaged for 10 consecutive bursts at each of 3 time points: control (5 minutes prior to perfusion), Acute, and after 30 minutes of modulation. There was no significant difference between control and acute conditions, but a significant increase from acute to 30 minutes (paired t-test, p<0.01). N=8. **D.** Interburst Interval decreases sharply in 5HT alone, and in 5HT+DA (p<0.05 for each).

**Figure 6.**
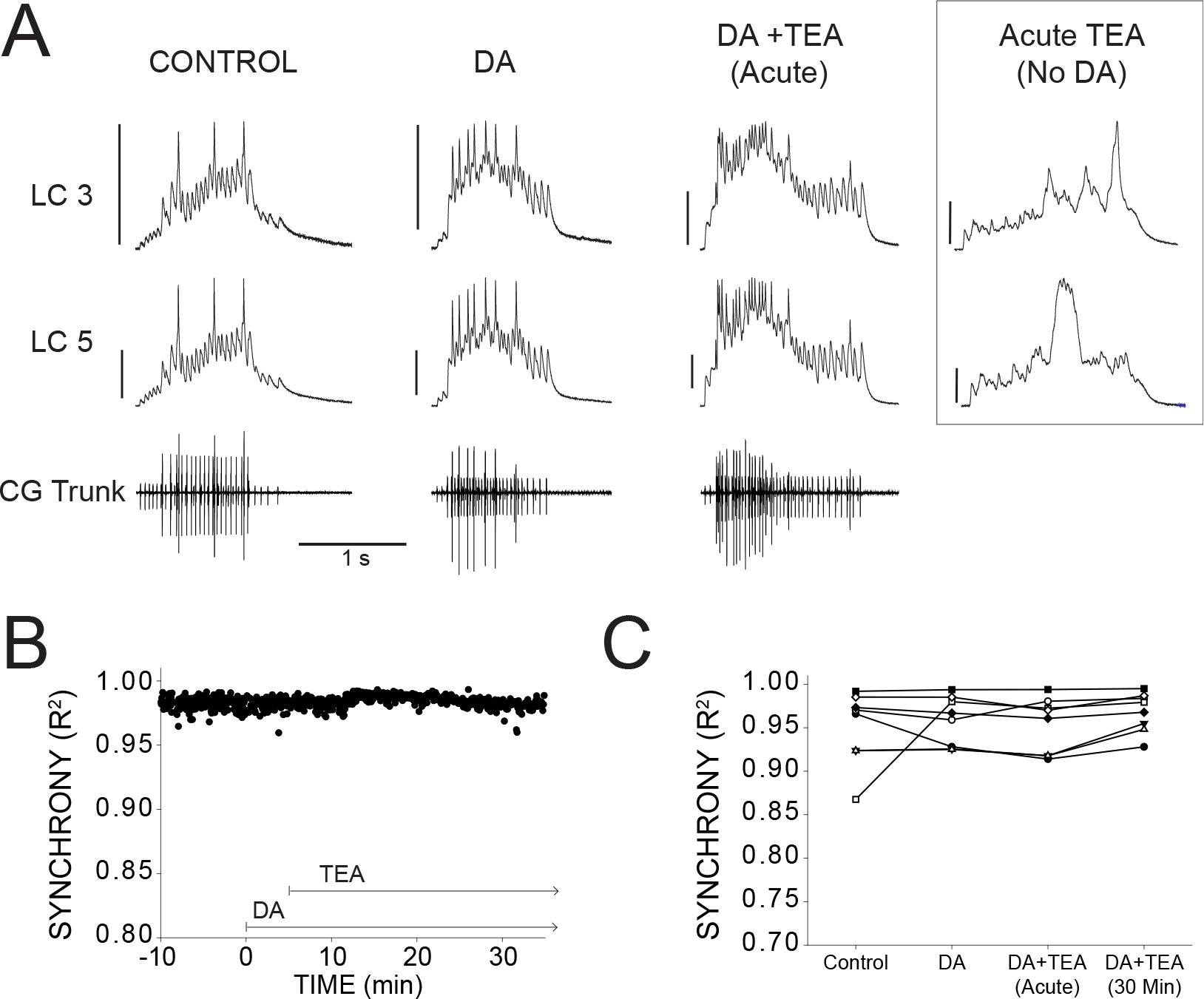
DA prevents desynchronization when co-applied with TEA. **A.** Representative traces for a single preparation in control, DA alone, and DA+TEA. Traces in the box at the far right illustrate acute desynchronization in TEA in the absence of DA (separate preparation). Scale bars = 10 mV and 1 second. **B.** R^2^ was calculated for every burst across a full experiment. Scatterplot shows throughout 10 minutes of control activity followed by 5 minutes of DA exposure, followed by 30 minutes in DA and TEA. **C.** R^2^ was averaged for 10 consecutive bursts at each of 4 time points: control (5 minutes prior to perfusion), after 5 minutes in DA, at the mean time point for desynchronization in TEA observed previously, and after 30 minutes exposure to the DA+TEA saline. No significant differences were detected across groups. N=8 preparations.

### Effects of DA and 5HT on crustacean motor neurons

Both 5HT and DA have been extensively studied in crustacean motor neurons, particularly those of the crustacean stomatogastric ganglion (STG). In particular, 5HT is known to reduce the conductance of both transient and persistent components of a calcium-dependent K^+^ current (I_KCa_) in STG cells, an effect that is mimicked by the application of tetraethylammonium [TEA] (*Kiehn and Harris-Warrick, 1992b*). In addition, 5HT enhances a slow voltage-dependent Ca^2+^ current (I_Ca_) in STG cells (*Kiehn and Harris-Warrick, 1992a; Zhang and Harris-Warrick, 1995*). These same I_KCa_ and I_Ca_ currents are known to be present in LCs of the cardiac network, and demonstrated to vary over a wide range of conductance magnitudes (*Ransdell et al., 2013a,b, 2012*). Furthermore, treatment of LCs with TEA results in an increase in excitability and a loss of network synchrony amongst LCs (*Ransdell et al., 2013a; Lane et al., 2016*). In addition, 5HT has only been reported to have very weak or no effect at all on electrical coupling in STG neurons *Johnson and Harris-Warrick, 1990*). These known features of serotonergic modulation in other crustacean motor neurons are therefore quite consistent with our data in this study. Specifically, 5HT causes an increase in excitability of LCs, as well as causes network desynchronization. We suggest that 5HT modulation has distinct effects on each LC within a ganglion due to the underlying variability of K^+^ and Ca^2+^ currents in each cell, and yet because 5HT has no effect on coupling this results in distinct hyperexcitable outputs from each cell that manifests as loss of synchronous activity.

DA has distinct and widespread effects on STG cells, both in terms of output and subcellular targets. DA is known to target the transient potassium current I_A_ (*Kloppenburg et al., 1999; Zhang et al., 2010*) as well as I_H_, I_NaP_, and I_Ca_ in STG cells (*Harris-Warrick, 2010*). The influence of DA on each conductance type is dependent on cell type (*Harris-Warrick, 2010; Zhang et al., 2010*) as well as the concentration and application time course of the modulator (*Rodgers et al., 2011, 2013*). Furthermore, DA has been shown to both increase and decrease electrical coupling among STG cells in a cell-type-specific manner *Johnson et al., 1993*). While I_H_ is not present in crab cardiac LCs, I_A_, I_NaP_, and I_Ca_ have all been identified and shown to vary over a wide range of conductance magnitudes in these cells (*Ransdell et al., 2013b, 2012*). Pharmacological blockade of I_A_ in LCs also has been shown to induce hyperexcitability and loss of synchrony in cardiac network output (*Ransdell et al., 2012*). Therefore, our prediction was that we should see similar loss of synchrony with DA application in this study. This was clearly not the case - while our data demonstrate an increase in excitability of LCs exposed to DA, they never experience any decrease in synchrony. In fact, synchrony was almost always enhanced as a result of DA application. It was only when we determine that DA also directly increases electrical coupling that we could propose a mechanism for the maintenance of synchrony despite the targeting of variable underlying conductances across LCs in DA. Namely, that DA’s influence on coupling prevents desynchronization and is effectively protective against the potentially variable impacts of modulation across LCs within the network. This hypothesis was tested by co-applying both 5HT and DA as well as TEA and DA, and the fact that neither of these conditions resulting in loss of synchrony is consistent with the hypothesis of a protective effect of increased coupling via DA in the cardiac network.

### Co-application of DA and 5HT has additive complementary effects on network output

While we directly tested the prediction of DA’s ability to preserve network synchrony in the face of divergent effects on variable LCs within a ganglion through co-application with both TEA and 5HT, we had no prediction as to what the combined influence of 5HT and DA would be on the cardiac network output overall. It is well established that individual biogenic amine modulators have distinct effects on both the CG (*Miller et al., 1984; Cruz-Bermudez and Marder, 2007*) and the STG (*Flamm and Harris-Warrick, 1986b,a*). So much so that distinct output modes can be attributed to DA, 5HT, and octopamine individually (*Flamm and Harris-Warrick, 1986b*), and is thought to underlie the diversity of circuit reorganization and outputs that has been demonstrated in these networks previously. Furthermore, few co-modulation experiments have been performed that specifically determine whether modulator effects will enhance, occlude, or have distinct effects from one another. Therefore, we did not anticipate that DA and 5HT would have largely complementary, and perhaps even synergistic effects on the ganglion. Our data demonstrate when these modulators were coapplied, the network output took on the hallmarks of 5HT modulation - including double-bursting of the LCs, enhanced excitability, and changes in spike frequency and bursting output - yet were protected from the loss of synchronous activity that accompanies 5HT-only application. These results suggest that DA and 5HT in these cells have distinct signaling pathways, subcellular targets, and ultimately result in a more linear combination of modulator effects than occlusion or saturation (*Li et al., 2018*). It remains to be seen whether DA interacts in this fashion with other modulators, including neuropeptides (*Miller and Sullivan, 1981; Cruz-Bermudez and Marder, 2007*). However, our results indicate that DA acts to ensure robust activity through the coordinated, synchronous action of all 5 LC motor neurons.

Only three pairs of axons provide extrinsic innervation of the CG - one pair is dopaminergic, the others are cholinergic and GABAergic (*Delgado et al., 2000; Cooke, 2002*). The dopaminergic fibers have many synaptic connections on anterior LCs, their neuropil (in the vicinity of the sites of electrical coupling), and the posterior of the ganglion near the pacemaker cells (*Fort et al., 2004*). This projection pattern is well situated to provide a potential means for direct delivery of DA to the CG over the immediate time scales of neuromodulator release, in addition to hormonal exposure to DA, through fibers that are rapidly responsive to physiologically relevant stimuli (*Maynard, 1953; Guirguis and Wilkens, 1995; Jury and Watson, 2000; Fort et al., 2004*). This would potentially allow for feedback transmitted through dopaminergic fibers to rapidly “prime” the network for subsequent modulation delivered hormonally, thus preventing desynchronization as a result.

### Conclusion: Neuromodulation, Electrical Coupling, and Robustness of Network Output

While neuromodulation is largely a phenomenon associated with network plasticity and changes in network output, a more recent appreciation of the additional role of neuromodulation in network robustness has begun to emerge. It has been proposed that the somewhat diffuse and often opposing actions of modulators on different components of the same circuit may act to stabilize the modulated state of networks, preventing “overmodulation” that could render a network nonfunctional or pathological (*Harris-Warrick, 2010; Marder, 2012*). For example, 5HT increases the set of intrinsic and synaptic current magnitude combinations that can give rise to specific behaviors such as bursting in a half center oscillator (*Grashow et al., 2009*), ensuring a larger parameter space over which appropriate output can be generated in the context of multiple modulation and hence increasing robustness of output. More recently it has been shown that neuromodulation can compensate for temperature-induced loss of motor pattern output in the STG (*Städele et al., 2015*), thus acting to preserve appropriate network output. Electrical coupling itself, as well as modulation of coupling interactions, also play a role in circuit robustness, allowing for distributed network solutions to maximize output consistency under different conditions or with variable underlying physiology of network constituents (*Kepler et al., 1990; Gutierrez and Marder, 2013; Gutierrez et al., 2013; Marder et al., 2017*).

Modulation of coupling via DA may have convergent effects on such robustness across different taxa, as modulation of electrical synapses by DA is known to be involved in many circuits and species. DA directly modulates gap junctional conductance and network activity in horizontal cells (*Piccolino et al., 1984; He et al., 2000*), AII amacrine cells (*Kothmann et al., 2009*), and rod cells *Jin et al., 2015*) of the retina. DA modulation of coupling conductance is also involved in sensorimotor function during copulation in C. elegans (*Correa et al., 2015*). Furthermore DA has been shown capable of modulating the electrical component of the mixed electrical-chemical synapses of auditory afferents onto the fish Mauthner cell (*Pereda et al., 1992; Cachope et al., 2007; Cachope and Pereda, 2012*). While electrical coupling is known to support synchronized activity in many systems, and DA known to modulate such coupling across taxa, to our knowledge this is the first study that directly implicates the neuromodulation of gap junctional conductance in directly counteracting desynchronizing perturbations to enhance network robustness. Furthermore, DA can do so in this system and maintain the ability of other modulators, in this case 5HT, to alter output in an independent fashion that is protected from potentially detrimental or destabilizing alterations to coordinated network activity. This expands the understanding of how neuromodulators can interact to ensure appropriate flexibility in circuits to respond to changes in sensory feedback and produce appropriate context-specific output, but ensure that overall robustness of the network is maintained within an acceptable parameter space.

## Methods and Materials

### Animals

Adult male Jonah crabs, Cancer borealis, were purchased and shipped overnight from The Fresh Lobster Company (Gloucester, MA). Crabs were maintained in artificial seawater at 12°C until used. Crabs were anesthetized by keeping them on ice for 30 minutes prior to dissection. The complete cardiac ganglion was dissected from the animal and pinned out in a Sylgard-lined petri dish in chilled physiological saline (440mM NaCl, 26mM MgCl2,13mM CaCl2,11mM KCl, and 10mM HEPES, pH 7.4-7.5,12°C). Chemicals were obtained from Fisher Scientific unless otherwise noted.

### Electrophysiology

The CG network is comprised of 9 cells: 4 Small Cell (SC) pacemaker interneurons which give simultaneous excitatory input to 5 Large Cell (LC) motor neurons. Stainless steel pin electrodes were connected to differential AC Amplifier (A-M Systems model 1700) for extracellular recording. One pin was placed inside a petroleum jelly well built around the ganglionic trunk and the other placed in the bath outside the well. The ganglionic trunk contains axons from all 9 cells, and thus serves to monitor the spiking output of the entire network. During normal activity, LCs produce consistent levels of rhythmic output and pairs of LCs show nearly identical voltage waveforms. LC and pacemaker spikes are easily distinguishable by their relative amplitudes, and by intracellular potential changes in the LCs.

LC somata are easily visible within the nerve and can be individually desheathed for intracellular sharp electrode recordings. Intracellular sharp electrodes containing 3M KCl (8-25 megaOhm) were used to simultaneously monitor the voltage activity in the somata of two anterior LCs. All paired intracellular recordings were from LC3 and either LC4 or LC5. Amplifiers from Axon Instruments were used (AxoClamp 900A, MultiClamp 700B, AxoClamp 2B). Current clamp protocols were created and run using Clampex 10.3 software (Molecular Devices). Electrical coupling in the intact network was measured by hyperpolarizing current injection (1-6 nA) when LCs reached resting membrane potential between bursts. Cells were injected one at a time while measuring voltage changes in both cells. Coupling coefficients were calculated as the ratio: (dV_coupled cell_ / dV_InjectedCell_). For both amines, the coupling coefficient at control was compared to the measured value after 15 minutes of modulation. Changes in coupling coefficient could ultimately be influenced by two fundamentally different mechanisms: altered conductance of the non-junctional membrane, or modification of gap junctional conductance. LC3 and LC5 are coupled to one another and the rest of the network distal to the site of our recordings, making calculation of coupling conductance between these two cells problematic. Our reduced preparation of the thread ligatured LC4 LC5 branch provides an electrotonically compact two-cell preparation in which we could calculate coupling conductance (*Bennett, 1966*).

### Modulator application

Serotonin (5HT), dopamine (DA), and tetraethylammonium (TEA) were obtained from Tocris Biosciences. Whenever present, concentrations were as follows: 10^−6^ M5HT, 10^−5^ M DA, 25mMTEA. All perfusions occurred at a rate of approximately 2 ml/min. Solutions were pre-chilled to maintain a constant temperature of 12°C in the dish. LC variability can be exploited by applying the K+ channel blocker TEA exclusively to anterior LCs to desynchronize their activity (Lane et al. 2016). To test whether neuromodulation could desynchronize LC activity, we considered perfusion of neuromodulators over the entire network to be a more biologically relevant challenge to test synchrony. There is no reason to suspect any biological conditions in which only the anterior LCs would be exposed to modulators. In vivo, the entire CG is exposed to both serotonin (5HT) and dopamine (DA) as hormonal modulators released from the pericardial organ or other neurohormonal sites (*Cooke, 2002; Fort et al., 2004; Maynard, 1953*). In addition, a pair of extrinsic dopaminergic fibers innervates the CG from the thoracic ganglion and forms abundant synaptic contacts in both the anterior and posterior regions of the ganglion which provides a rapid and direct route for dopaminergic modulation (*Cooke, 2002; Fort et al., 2004*). Baseline activity was measured during a sham perfusion of physiological saline before the source of the perfusion was switched to saline containing neuromodulators and/or channel blockers. Simultaneous intracellular recordings monitored somatic burst potentials from multiple anterior LCs, and an extracellular well on the ganglionic trunk monitored activity of the entire network.

In some experiments, a petroleum jelly well was used to protect the posterior end of the ganglion from the perfusate to selectively expose the anterior LCs to 5HT, DA, or DA+TEA. For experiments in which TEA was applied only to the anterior portion of the network, whole-network modulation was achieved by adding DA to the posterior end of the ganglion by pipette at the same time the DA perfusion began.

### Data Analysis

Intracellular burst waveforms were considered to begin with the first EPSP from pacemaker activity and ended upon return to resting membrane potential. Recordings were analyzed using Clampfit 10.3 (Molecular Devices) and Spike 2.7 (CED, Cambridge, UK) software. Statistical analyses were performed using Sigmaplot 11.0. All data are expressed as mean ± SD unless otherwise stated. R-values (and R^2^) were obtained by Pearson correlation tests. Changes in burst characteristics, coupling coefficients, coupling conductance, and synchrony were analyzed with paired t-tests when data was normally distributed, or signed rank tests in the case of non-normality. All comparisons for measures of network output between control and modulation are after 15 minutes of modulation. Each burst characteristic quantified was averaged for 10 consecutive cycles at control and at 15 minutes.

The sample sizes to compare waveform synchrony were calculated with power analyses based on data reported in two previous studies with highly similar experimental manipulations of exposure of LCs to compounds that cause loss of synchrony (*Ransdell et al., 2013a; Lane et al., 2016*), which yielded target sample size of N=6-8 to yield a power of 0.8 or higher. Sample sizes for changes in network output following modulator exposure were based on similar data in our previous work (*Ransdell et al., 2012*), and yielded target sample sizes of N=5-6 to achieve a power of 0.8. Power analyses were conducted based on the use of paired t-tests to analyze the data. However, when data were not normally distributed we used Wilcoxon signed rank tests. All sample sizes used in our studies are reported in Figure Legends and/or in the Results section when significance values are reported.

## Acknowledgments

We would like to thank Dr. Joseph Santin for feedback on the manuscript.

## References

Ball JM, Franklin CC, Tobin AE, Schulz DJ, Nair SS. Coregulation of ion channel conductances preserves output in a computational model of a crustacean cardiac motor neuron. Journal of Neuroscience. 2010 jun; 30(25):8637–8649. http://www.jneurosci.org/cgi/doi/10.1523/JNEUROSCI.6435-09.2010, doi: 10.1523/JNEUROSCI.6435-09.2010.

Bargmann CI. Beyond the connectome: How neuromodulators shape neural circuits. BioEssays. 2012; 34(6):458–465. doi: 10.1002/bies.201100185.

Bennett MVL. Physiology of electrotonic junctions. Annals of the New York Academy of Sciences. 1966 jul; 137(2):509–539. doi: 10.1111/j.1749-6632.1966.tb50178.x.

Cachope R, Mackie K, Triller A, O’Brien J, Pereda AE. Potentiation of Electrical and Chemical Synaptic Transmission Mediated by Endocannabinoids. Neuron. 2007; 56(6):1034–1047. doi: 10.1016/j.neuron.2007.11.014.

Cachope R, Pereda AE, Two independent forms of activity-dependent potentiation regulate electrical transmission at mixed synapses on the Mauthner cell; 2012. doi: 10.1016/j.brainres.2012.05.059.

Calabrese RL, Norris BJ, Wenning A, Wright TM. Coping with variability in small neuronal networks. In: Integrative and Comparative Biology, vol. 51 Oxford University Press; 2011. p. 845–855. doi: 10.1093/icb/icr074.

Cooke IM, Reliable, responsive pacemaking and pattern generation with minimal cell numbers: The crustacean cardiac ganglion; 2002. doi: 10.2307/1543649.

Correa PA, Gruninger T, Garcia LR. DOP-2 D2-Like Receptor Regulates UNC-7 Innexins to Attenuate Recurrent Sensory Motor Neurons during C. elegans Copulation. Journal of Neuroscience. 2015; 35(27):9990–10004. http://www.jneurosci.org/cgi/doi/10.1523/JNEUROSCI.0940-15.2015, doi: 10.1523/JNEUROSCI.0940-15.2015.

Cruz-Bermudez ND, Marder E. Multiple modulators act on the cardiac ganglion of the crab, Cancer bore alis. Journal of Experimental Biology. 2007 aug; 210(16):2873–2884. http://jeb.biologists.org/cgi/doi/10.1242/jeb.002949, doi: 10.1242/jeb.002949.

Daur N, Nadim F, Bucher D, The complexity of small circuits: the stomatogastric nervous system. Elsevier Ltd; 2016. doi: 10.1016/j.conb.2016.07.005.

Delgado JY, Oyola E, Miller MW. Localization of GABA- and glutamate-like immunoreactivity in the cardiac ganglion of the lobster Panulirus argus. Journal of Neurocytology. 2000; 29(8):605–619. http://link.springer.com/10.1023/A:1011080304472, doi: 10.1023/A:1011080304472.

Flamm RE, Harris-Warrick RM. Aminergic modulation in lobster stomatogastric ganglion.I. Effects on motor pattern and activity of neurons within the pyloric circuit. Journal of Neurophysiology. 1986; 55(5):847–865. doi: 10.1152/jn.1986.55.5.847.

Flamm RE, Harris-Warrick RM. Aminergic modulation in lobster stomatogastric ganglion. II. Target neurons of dopamine, octopamine, and serotonin within the pyloric circuit. Journal of neurophysiology. 1986; 55(5):866–881. doi: 10.1152/jn.1986.55.5.866.

Fort TJ, Brezina V, Miller MW. Modulation of an integrated central pattern generator-effector system: Dopaminergic regulation of cardiac activity in the blue crab Callinectes sapidus. Journal of Neurophysiology. 2004 dec; 92(6):3455–3470. http://jn.physiology.org/cgi/doi/10.1152/jn.00550.2004, doi: 10.1152/jn.00550.2004.

Goldman MS, Golowasch J, Marder E, Abbott LF. Global structure, robustness, and modulation of neuronal models. TheJournal of neuroscience: the official journal of the Society for Neuroscience. 2001; 21(14):5229–38. http://www.ncbi.nlm.nih.gov/pubmed/11438598, doi: 21/14/5229[pii].

Grashow R, Brookings T, Marder E. Reliable neuromodulation from circuits with variable underlying structure. Proceedings of the National Academy of Sciences. 2009; 106(28):11742–11746. http://www.pnas.org/lookup/doi/10.1073/pnas.0905614106, doi: 10.1073/pnas.0905614106.

Gruhn M, Guckenheimer J, Land B, Harris-Warrick RM. Dopamine modulation of two delayed rectifier potassium currents in a small neural network. Journal of Neurophysiology. 2005; 94(4):2888–900. http://www.ncbi.nlm.nih.gov/pubmed/16014791, doi: 10.1152/jn.00434.2005.

Guirguis MS, Wilkens JL. The role of the cardioregulatory nerves in mediating heart rate responses to locomotion, reduced stroke volume, and neurohormones in Homarus americanus. Biological Bulletin. 1995; 188(2):179–185. doi: 10.2307/1542083.

Gutierrez GJ, Marder E. Rectifying electrical synapses can affect the influence of synaptic modulation on output pattern robustness. Journal of Neuroscience. 2013 aug; 33(32):13238–13248. http://www.jneurosci.org/cgi/doi/10.1523/JNEUROSCI.0937-13.2013, doi: 10.1523/JNEUROSCI.0937-13.2013.

Gutierrez GJ, Marder E. Modulation of a single neuron has state- dependent actions on circuit dynamics. eNeuro. 2014; 1(December):1–12. doi: 10.1523/ENEURO.0009-14.

Gutierrez GJ, O’Leary T, Marder E. Multiple mechanisms switch an electrically coupled, synaptically inhibited neuron between competing rhythmic oscillators. Neuron. 2013 mar; 77(5):845–858. doi: 10.1016/j.neuron.2013.01.016.

Harris-Warrick. Checks and balances in neuromodulation. Frontiers in Behavioral Neuroscience. 2010; 4(July):1–9. http://journal.frontiersin.org/article/10.3389/fnbeh.2010.00047/abstract, doi: 10.3389/fnbeh.2010.00047.

Harris-Warrick RM, Neuromodulation and flexibility in Central Pattern Generator networks. Elsevier Ltd; 2011. doi: 10.1016/j.conb.2011.05.011.

He S, Weiler R, Vaney DI. Endogenous dopaminergic regulation of horizontal cell coupling in the mammalian retina. Journal of Comparative Neurology. 2000; 418(1):33–40. doi: 10.1002/(SICI)1096-9861(20000228)418:1<33::AID-CNE3>3.0.CO;2-J.

Jin NG, Chuang AZ, Masson PJ, Ribelayga CP. Rod electrical coupling is controlled by a circadian clock and dopamine in mouse retina. Journal of Physiology. 2015; 593(7):1597–1631. doi: 10.1113/jphysiol.2014.284919.

Johnson BR, Peck JH, Harris-Warrick RM. Amine modulation of electrical coupling in the pyloric network of the lobster stomatogastric ganglion. Journal of Comparative Physiology A. 1993; 172(6):715–732. doi: 10.1007/BF00195397.

Johnson BR, Harris-Warrick RM. Aminergic modulation of graded synaptic transmission in the lobster stomatogastric ganglion. The Journal of Neuroscience. 1990; 10(7):2066–76. http://www.ncbi.nlm.nih.gov/pubmed/2165519.

Johnson BR, Kloppenburg P, Harris-Warrick RM. Dopamine modulation of calcium currents in pyloric neurons of the lobster stomatogastric ganglion. Journal of Neurophysiology. 2003; 90(2):631–643. doi: 10.1152/jn.00037.2003.

Jury SH, Watson WH. Thermosensitivity of the lobster, Homarus americanus, as determined by cardiac assay. Biological Bulletin. 2000; 199(3):257–264. doi: 10.2307/1543182.

Kepler TB, Marder EVE, Abbott LF. The effect of electrical coupling on the frequency of model neuronal oscillators. Science. 1990 apr; 248(4951):83–85. doi: 10.1126/science.2321028.

Kiehn O, Harris-Warrick RM. 5-HT modulation of hyperpolarization-activated inward current and calcium-dependent outward current in a crustacean motor neuron. Journal of neurophysiology. 1992; 68(2):496–508.

Kiehn O, Harris-Warrick RM. Serotonergic stretch receptors induce plateau properties in a crustacean motor neuron by a dual-conductance mechanism. Journal of neurophysiology. 1992; 68(2):485–95. http://www.ncbi.nlm.nih.gov/pubmed/1527571.

Kloppenburg P, Levini RM, Harris-Warrick RM. Dopamine modulates two potassium currents and inhibits the intrinsic firing properties of an identified motor neuron in a central pattern generator network. Journal of Neurophysiology. 1999; 81:29–38.

Kothmann WW, Massey SC, O’Brien J. Dopamine-stimulated dephosphorylation of connexin 36 mediates aii amacrine cell uncoupling. Journal of Neuroscience. 2009 nov; 29(47):14903–14911. http://www.jneurosci.org/cgi/doi/10.1523/JNEUROSCI.3436-09.2009, doi: 10.1523/JNEUROSCI.3436-09.2009.

Lane BJ, Samarth P, Ransdell JL, Nair SS, Schulz DJ. Synergistic plasticity of intrinsic conductance and electrical coupling restores synchrony in an intact motor network. eLife. 2016; 5(AUGUST):1–23. doi: 10.7554/eLife.16879.

Li X, Bucher DM, Nadim F. Distinct co-modulation rules of synaptic and voltage-gated currents coordinates interactions of multiple neuromodulators. bioRxiv. 2018; https://www.biorxiv.org/content/early/2018/02/22/265694.full.pdf+html.

Marder E. Variability, compensation, and modulation in neurons and circuits. Proceedings of the National Academy of Sciences. 2011 sep; 108(Supplement_3):15542–15548. http://www.pnas.org/cgi/doi/10.1073/pnas.1010674108, doi: 10.1073/pnas.1010674108.

Marder E, Neuromodulation of neuronal circuits: Back to the future. Elsevier Inc.; 2012. doi: 10.1016/j.neuron.2012.09.010.

Marder E, Gutierrez GJ, Nusbaum MP. Complicating connectomes: Electrical coupling creates parallel pathways and degenerate circuit mechanisms. Developmental Neurobiology. 2017; 77(5):597–609. doi: 10.1002/dneu.22410.

Marder E, O’Leary T, Shruti S. Neuromodulation of circuits with variable parameters: Single neurons and small circuits reveal principles of state-dependent and robust neuromodulation. Annual Review of Neuroscience. 2014; 37(1):329–346. http://www.annualreviews.org/doi/10.1146/annurev-neuro-071013-013958, doi: 10.1146/annurev-neuro-071013-013958.

Maynard DM. Activity in a crustacean ganglion. I. Cardio-inhibition and accelaration in Panulirus argus. The Biological Bulletin. 1953; 104(2):156–170. http://www.biolbull.org/content/104/2/156.abstract.

Miller MW, Benson JA, Berlind A. Excitatory effects of dopamine on the cardiac ganglia of the crabs Portunus sanguinolentus and Podophthalmus vigil. Journal of Experimental Biology. 1984; 108(1). http://jeb.biologists.org/content/108/1/97.

Miller MW, Sullivan RE. Some effects of proctolin on the cardiac ganglion of the maine lobster, Homarus americanus. Journal of Neurobiology. 1981; 12(6):629–639. doi: 10.1002/neu.480120611.

Nadim F, Brezina V, Destexhe A, Linster C. State dependence of network output: Modeling and experiments. Journal of Neuroscience. 2008; 28(46):11806–11813. http://www.jneurosci.org/cgi/doi/10.1523/JNEUROSCI.3796-08.2008, doi: 10.1523/JNEUROSCI.3796-08.2008.

Nadim F, Bucher D. Neuromodulation of neurons and synapses. Current Opinion in Neurobiology. 2014; 29:48–56. http://dx.doi.org/10.1016/jxonb.2014.05.003, doi: 10.1016/j.conb.2014.05.003.

Peck JH, Nakanishi ST, Yaple R, Harris-Warrick RM. Amine modulation of the transient potassium current in identified cells of the lobster stomatogastric ganglion. Journal of Neurophysiology. 2001; 86(6):2957–65. http://www.ncbi.nlm.nih.gov/pubmed/11731552.

Pereda A, Triller A, Korn H, Faber DS. Dopamine enhances both electrotonic coupling and chemical excitatory postsynaptic potentials at mixed synapses. Proceedings of the National Academy of Sciences of the United States of America. 1992 dec; 89(24):12088–12092. doi: 10.1073/pnas.89.24.12088.

Piccolino M, Neyton J, Gerschenfeld HM. Decrease of gap junction permeability induced by dopamine and cyclic adenosine 3′:5′-monophosphate in horizontal cells of turtle retina. Journal of Neuroscience. 1984; 4(10):2477–88. doi: 10.1523/JNEUROSCI.04-10-02477.1984.

Ransdell JL, Nair SS, Schulz DJ. Rapid Homeostatic Plasticity of Intrinsic Excitability in a Central Pattern Generator Network Stabilizes Functional Neural Network Output. Journal of Neuroscience. 2012 jul; 32(28):9649–9658. http://www.jneurosci.org/cgi/doi/10.1523/JNEUROSCI.1945-12.2012, doi: 10.1523/JNEUROSCI.1945-12.2012.

Ransdell JL, Nair SS, Schulz DJ. Neurons within the same network independently achieve conserved output by differentially balancing variable conductance magnitudes. Journal of Neuroscience. 2013 jun; 33(24):9950–9956. http://www.jneurosci.org/cgi/doi/10.1523/JNEUROSCI.1095-13.2013, doi: 10.1523/JNEUROSCI.1095- 13.2013.

Ransdell JL, Temporal S, West NL, Leyrer ML, Schulz DJ. Characterization of inward currents and channels underlying burst activity in motoneurons of crab cardiac ganglion. Journal of Neurophysiology. 2013 jul; 110(1):42–54. http://jn.physiology.org/cgi/doi/10.1152/jn.00009.2013, doi: 10.1152/jn.00009.2013.

Rodgers EW, Krenz WDC, Baro DJ. Tonic Dopamine Induces Persistent Changes in the Transient Potassium Current through Translational Regulation. Journal of Neuroscience. 2011 sep; 31(37):13046–13056. http://www.jneurosci.org/cgi/doi/10.1523/JNEUROSCI.2194-11.2011, doi: 10.1523/JNEUROSCI.2194-11.2011.

Rodgers EW, Krenz WD, Jiang X, Li L, Baro DJ. Dopaminergic tone regulates transient potassium current maximal conductance through a translational mechanism requiring D1Rs, cAMP/PKA, Erk and mTOR. BMC neuroscience. 2013; 14:143. doi: 10.1186/1471-2202-14-143.

Städele C, Heigele S, Stein W. Neuromodulation to the Rescue: Compensation of Temperature-Induced Breakdown of Rhythmic Motor Patterns via Extrinsic Neuromodulatory Input. PLoS Biology. 2015; 13(9):e1002265. doi: 10.1371/journal.pbio.1002265.

Williams AH, Calkins A, O’Leary T, Symonds R, Marder E, Dickinson PS. The neuromuscular transform of the lobster cardiac system explains the opposing effects of a neuromodulator on muscle output. Journal of Neuroscience. 2013; 33(42):16565–16575. http://www.jneurosci.org/cgi/doi/10.1523/JNEUROSCI.2903-13.2013, doi: 10.1523/JNEUROSCI.2903-13.2013.

Zhang B, Harris-Warrick RM. Calcium-Dependent Plateau Potentials in a Crab Stomatogastric Ganglion Motor Neuron .1. Calcium Current and Its Modulation By Serotonin. Journal of Neurophysiology. 1995; 74(5):1929–1937.

Zhang H, Rodgers EW, Krenz WDC, Clark MC, Baro DJ. Cell specific dopamine modulation of the transient potassium current in the pyloric network by the canonical D1 receptor signal transduction cascade. Journal of Neurophysiology. 2010; 104:873–884. doi: 10.1152/jn.00195.2010.

